## Supplementary figures and images for "Transcriptomic comparison of two selective retinal cell ablation paradigms in zebrafish reveals shared and cell-specific regenerative responses"

### Supp1.tif

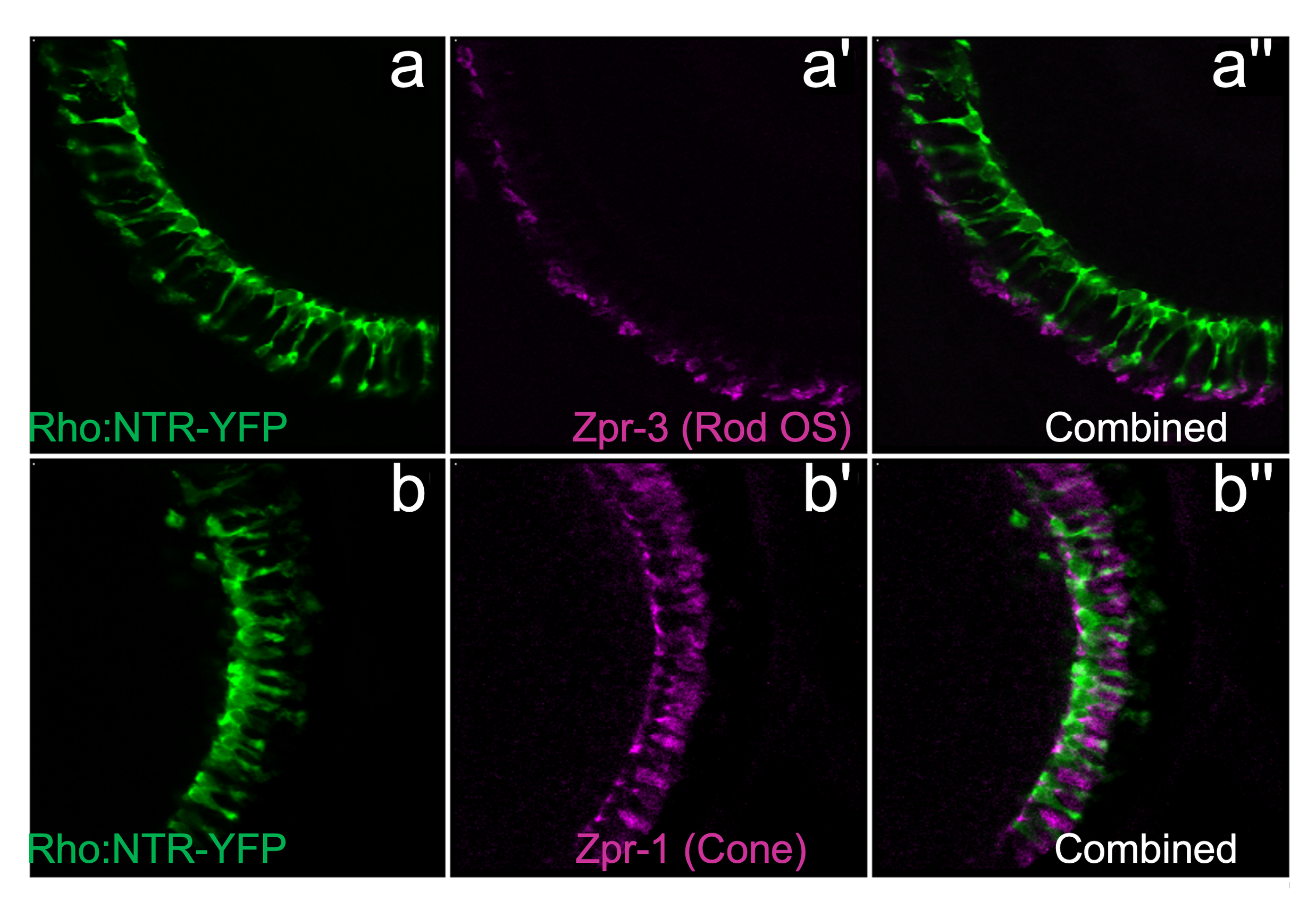

### Supp2.tif

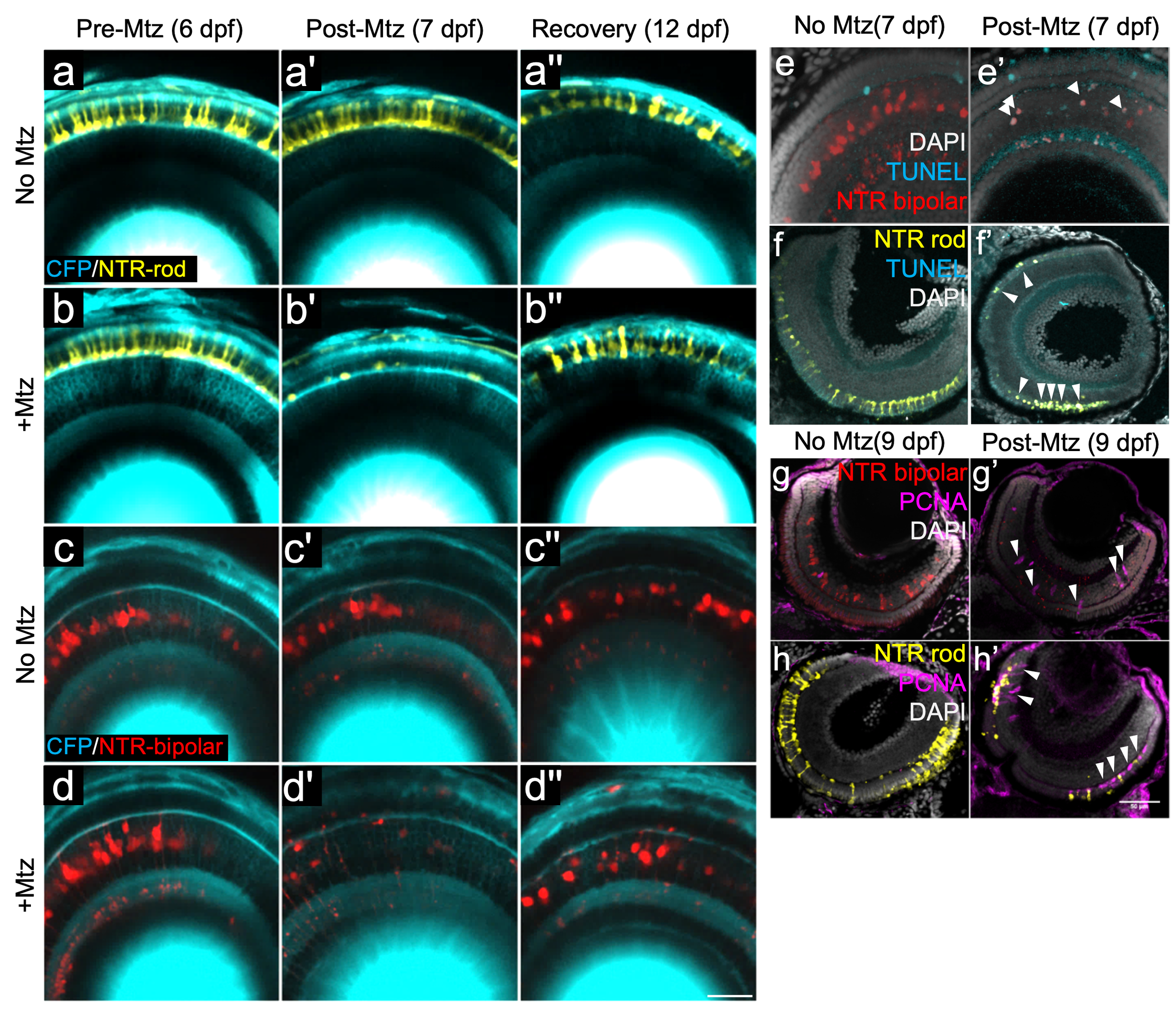

### Supp3.tif

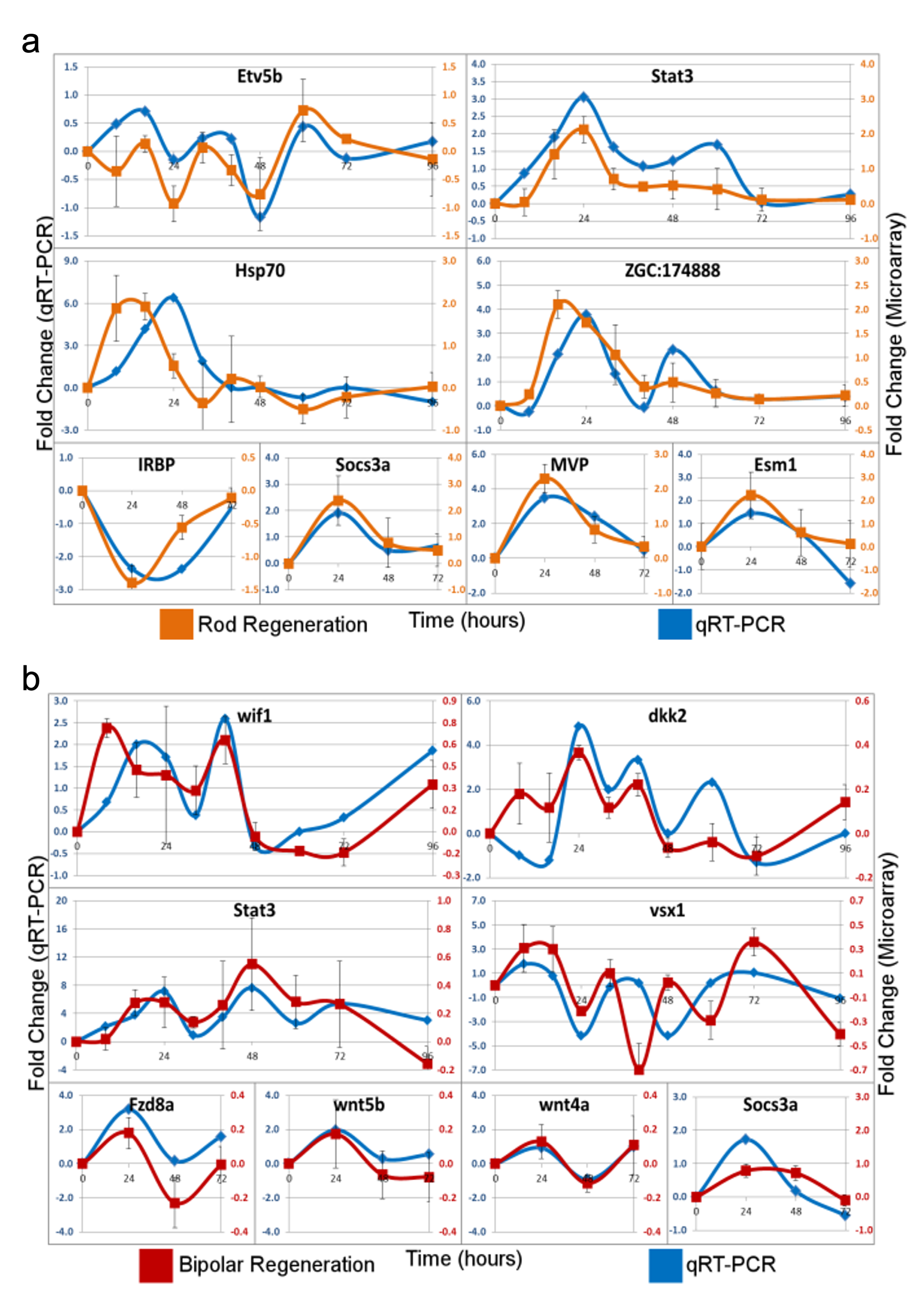

### Supp4.tif

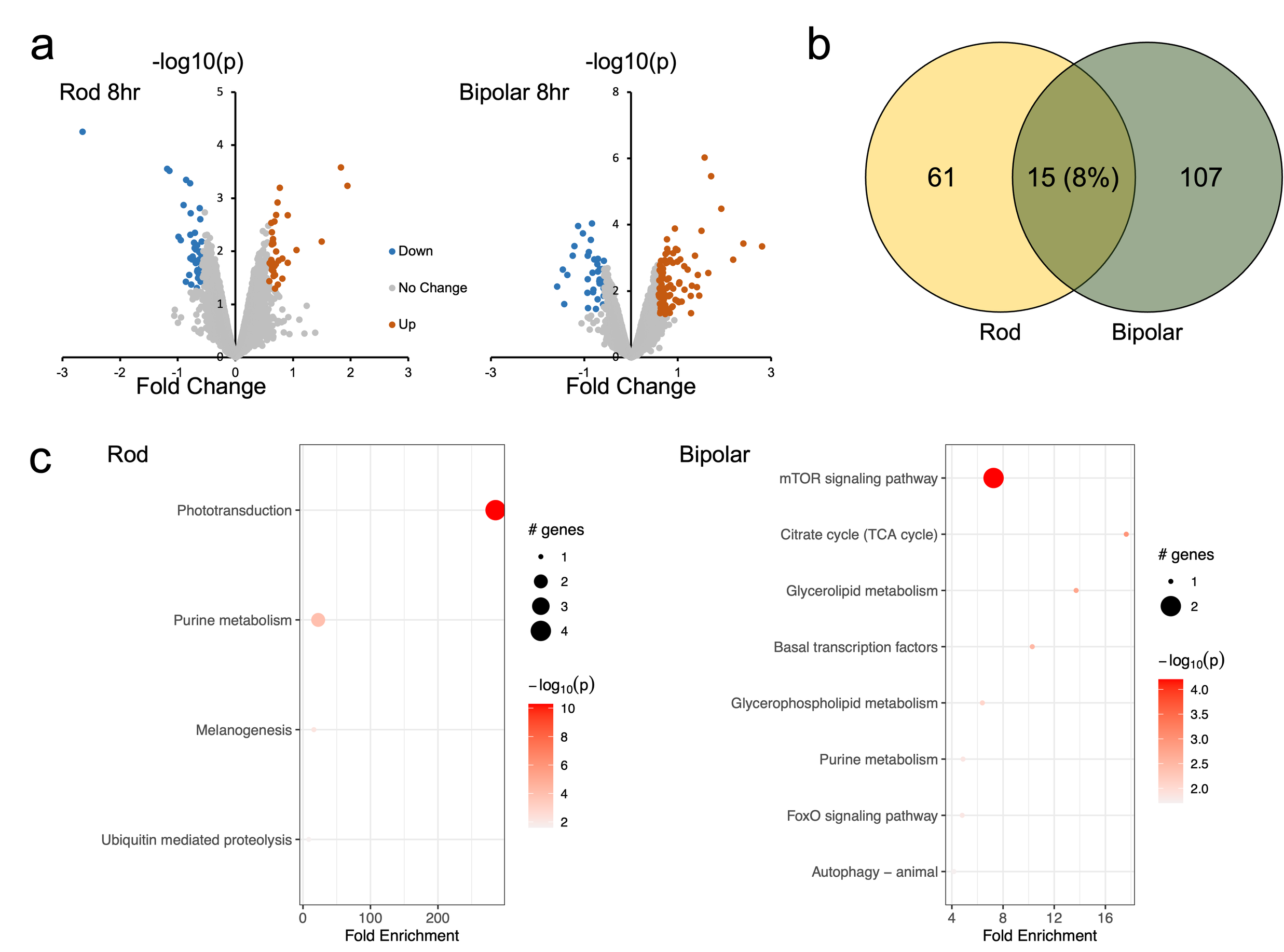

### Supp5.tif

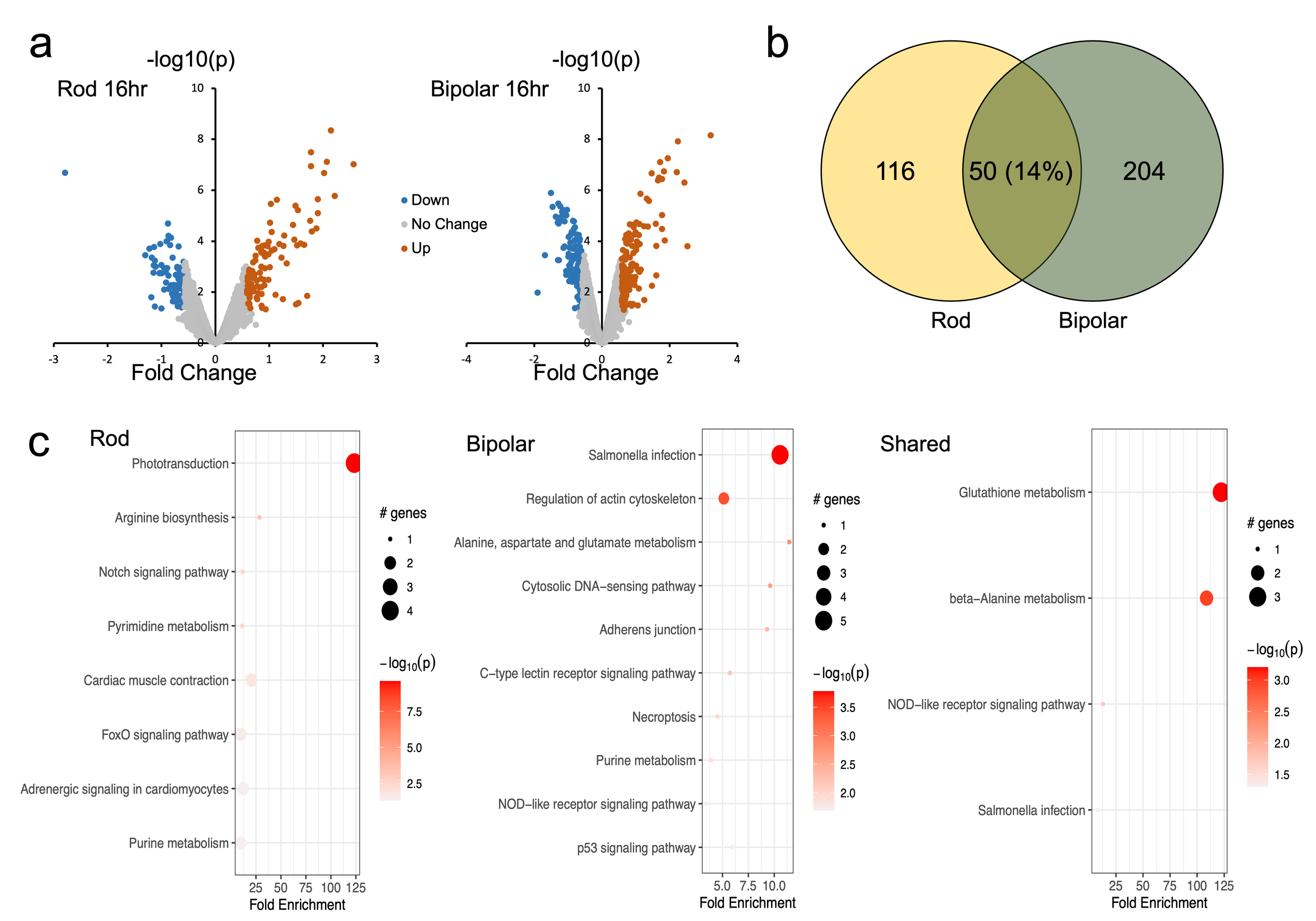

### Supp6.tif

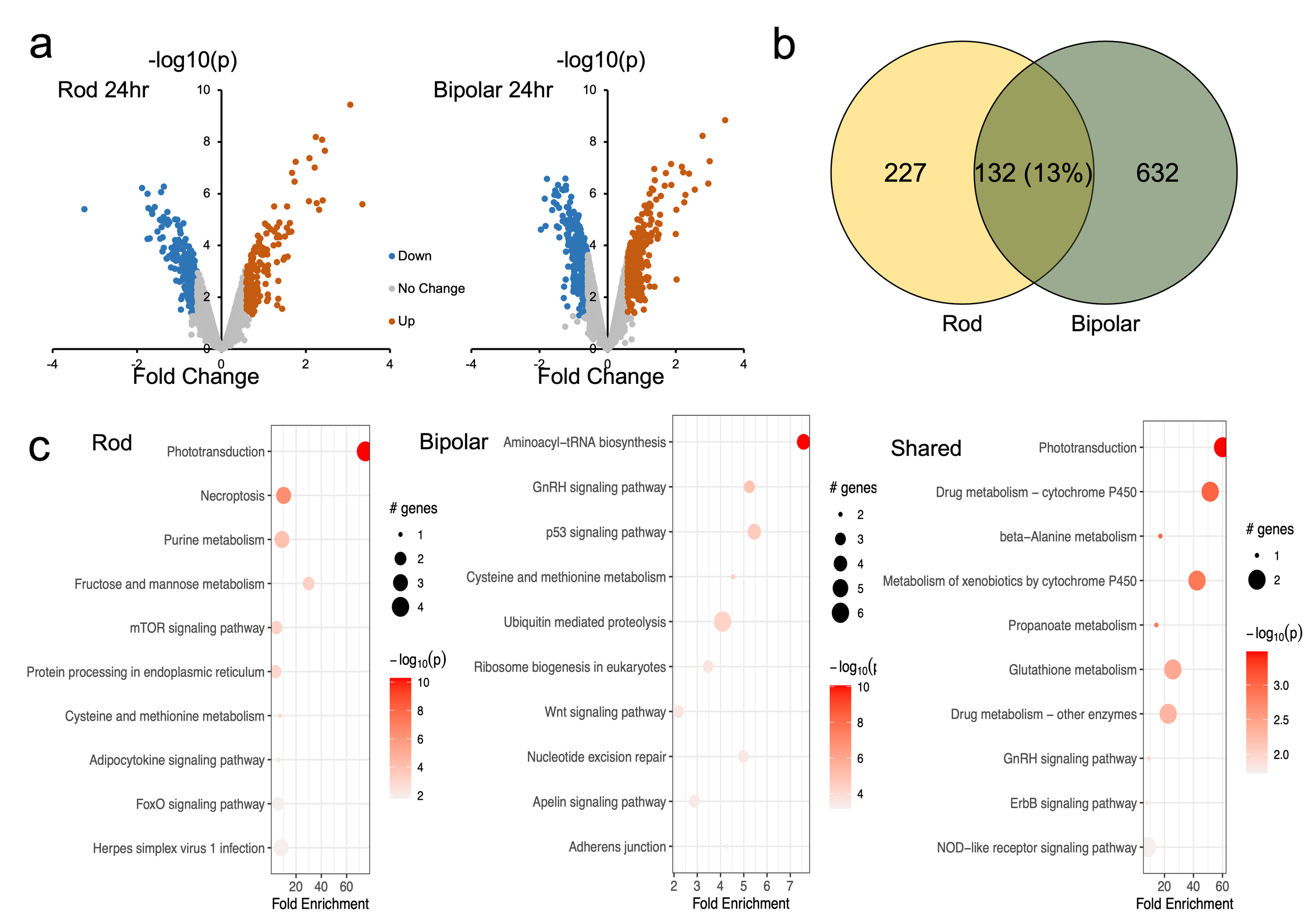

### Supp7.tif

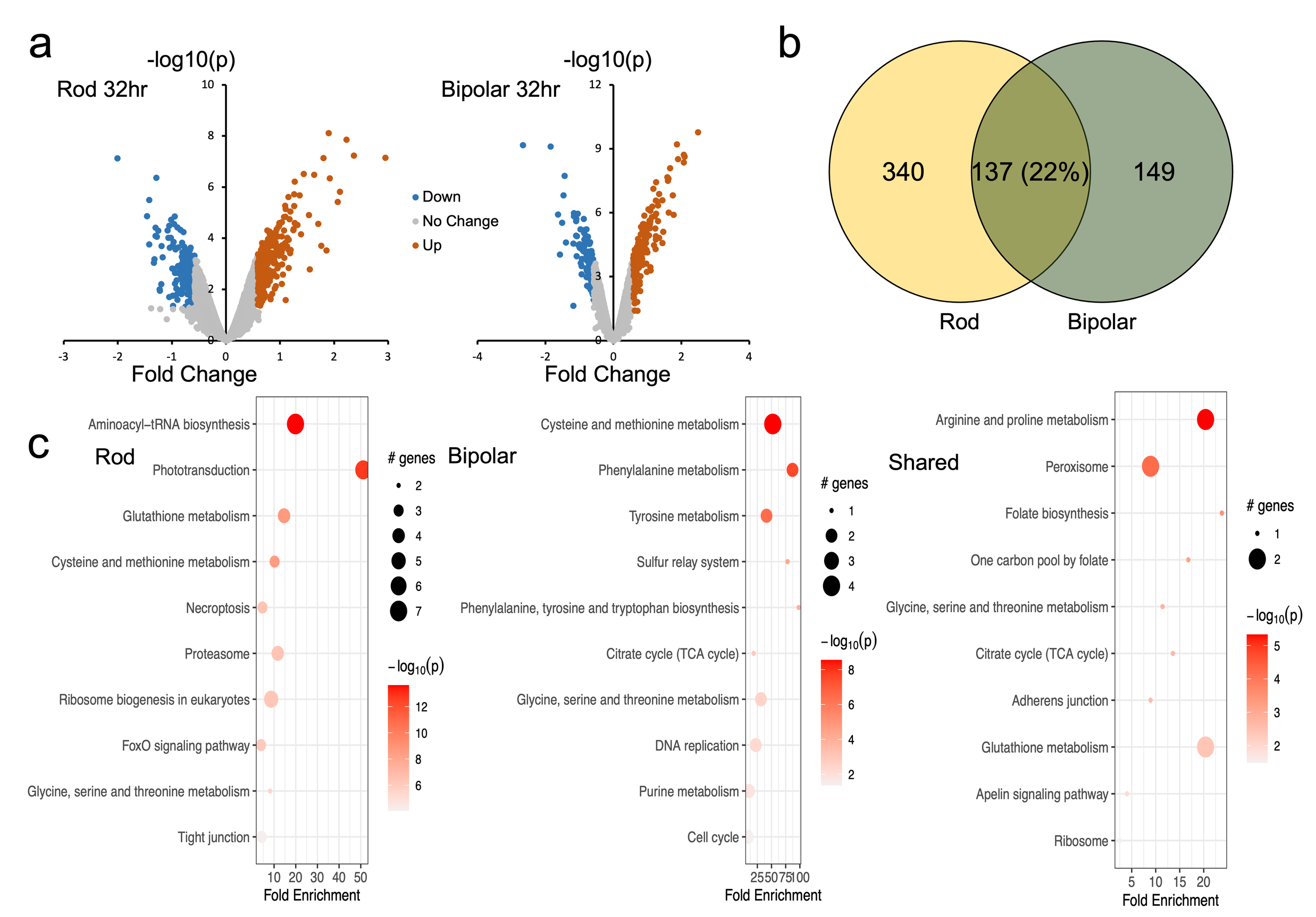

### Supp8.tif

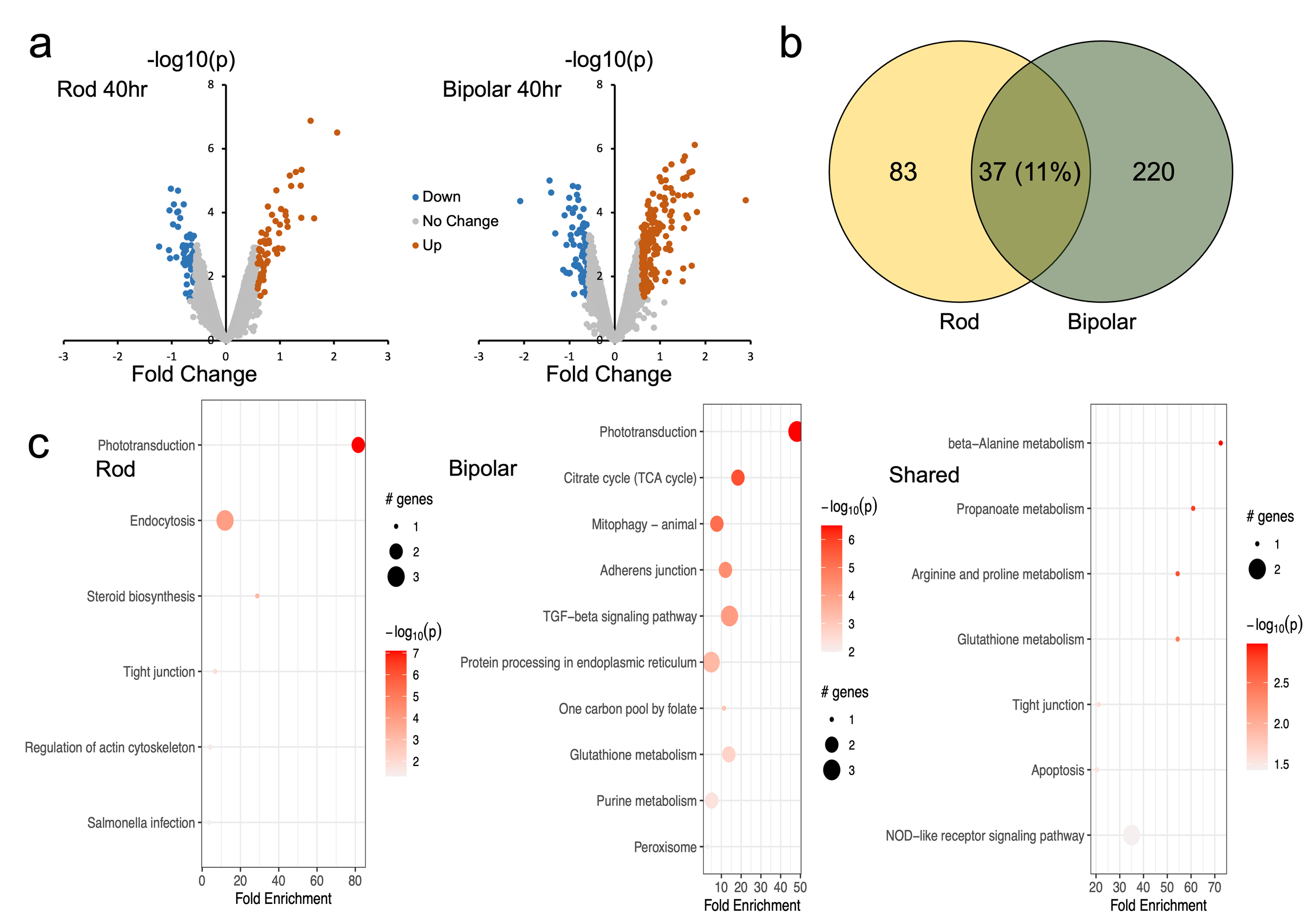

### Supp9.tif

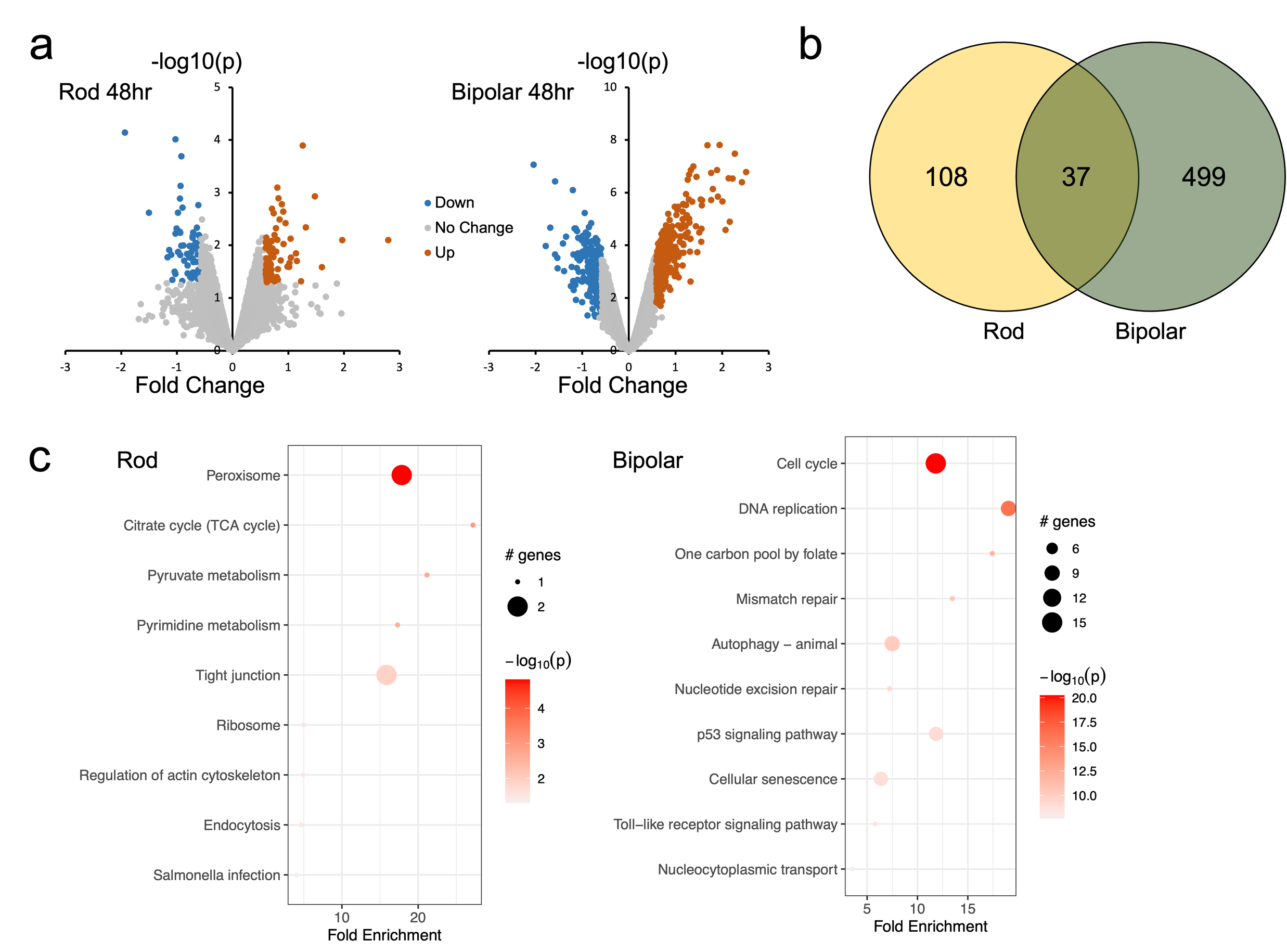

### Supp10.tif

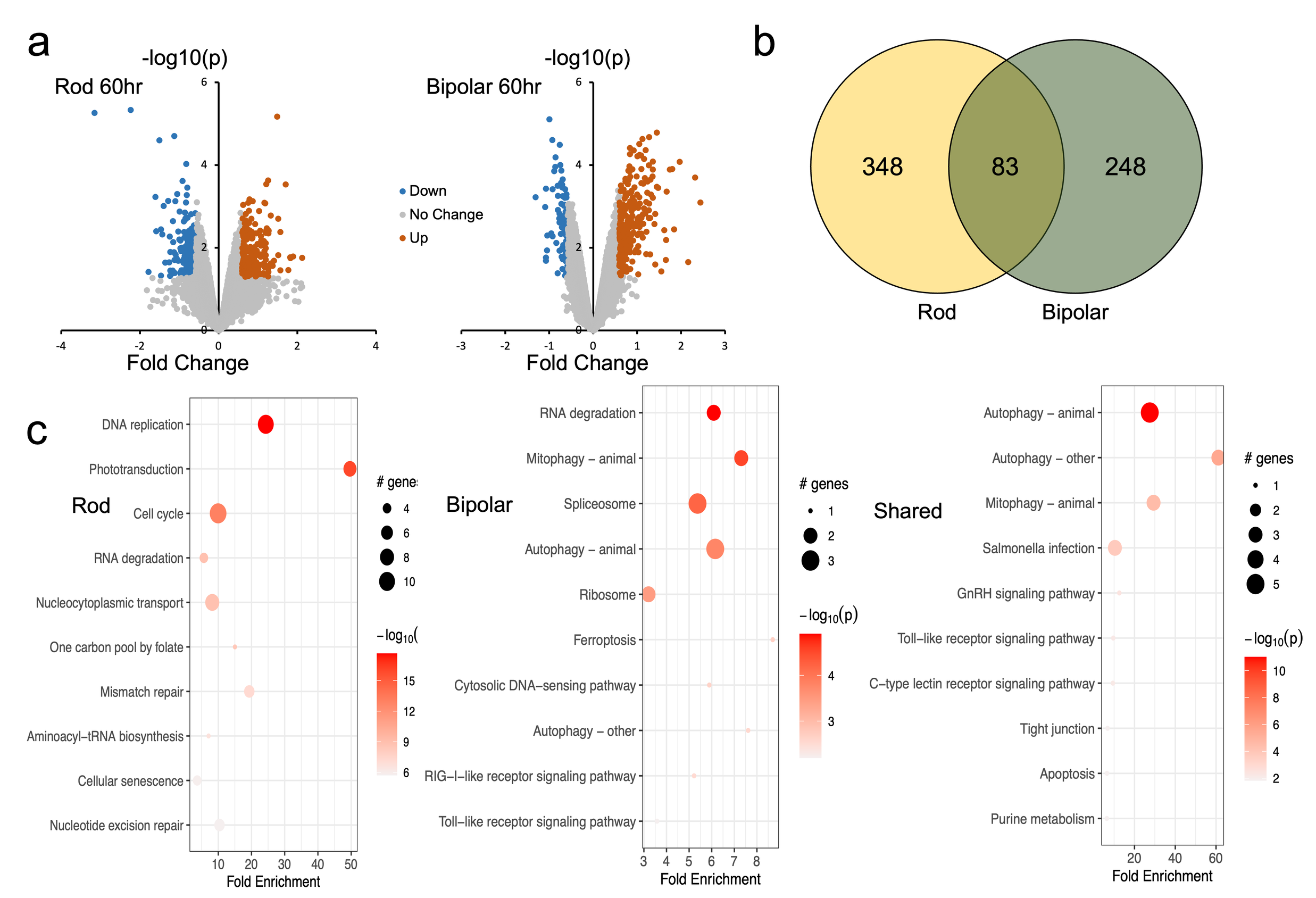

### Supp11.tif

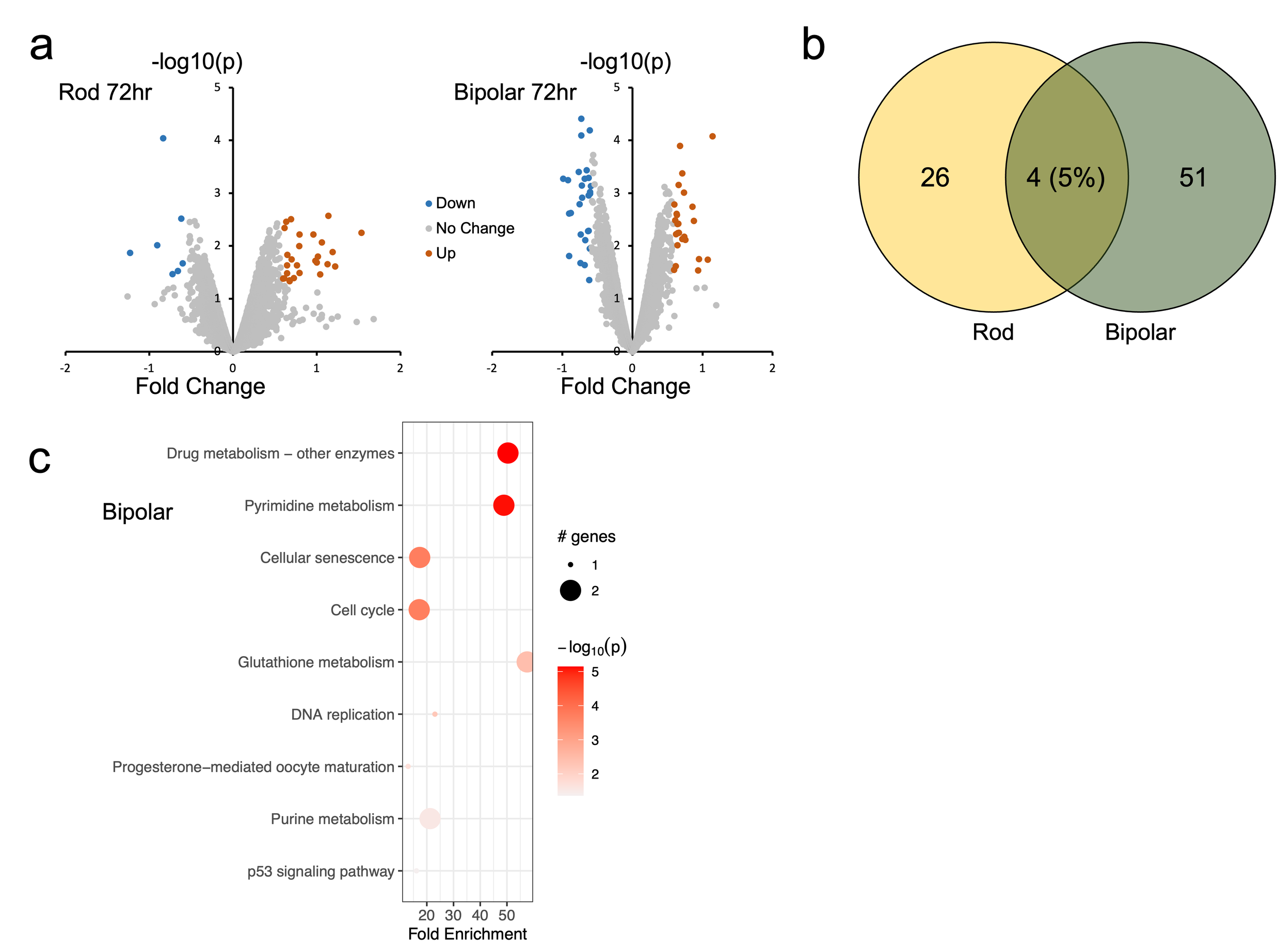

### Supp12.tif

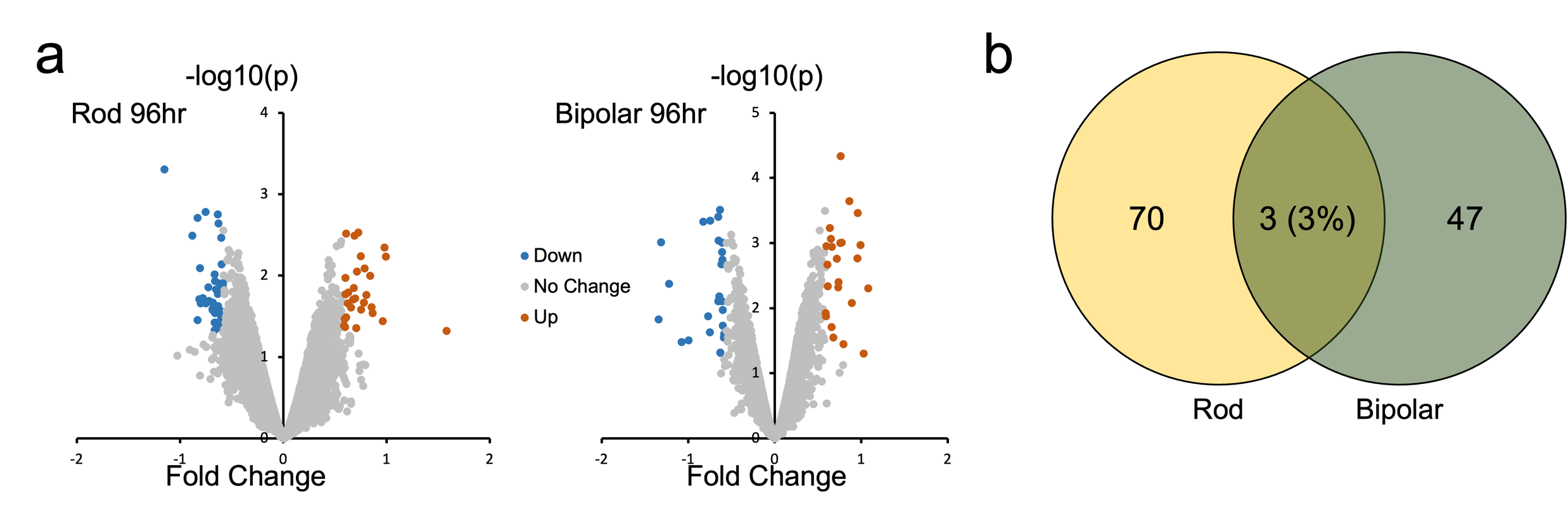

### Supp13.tif

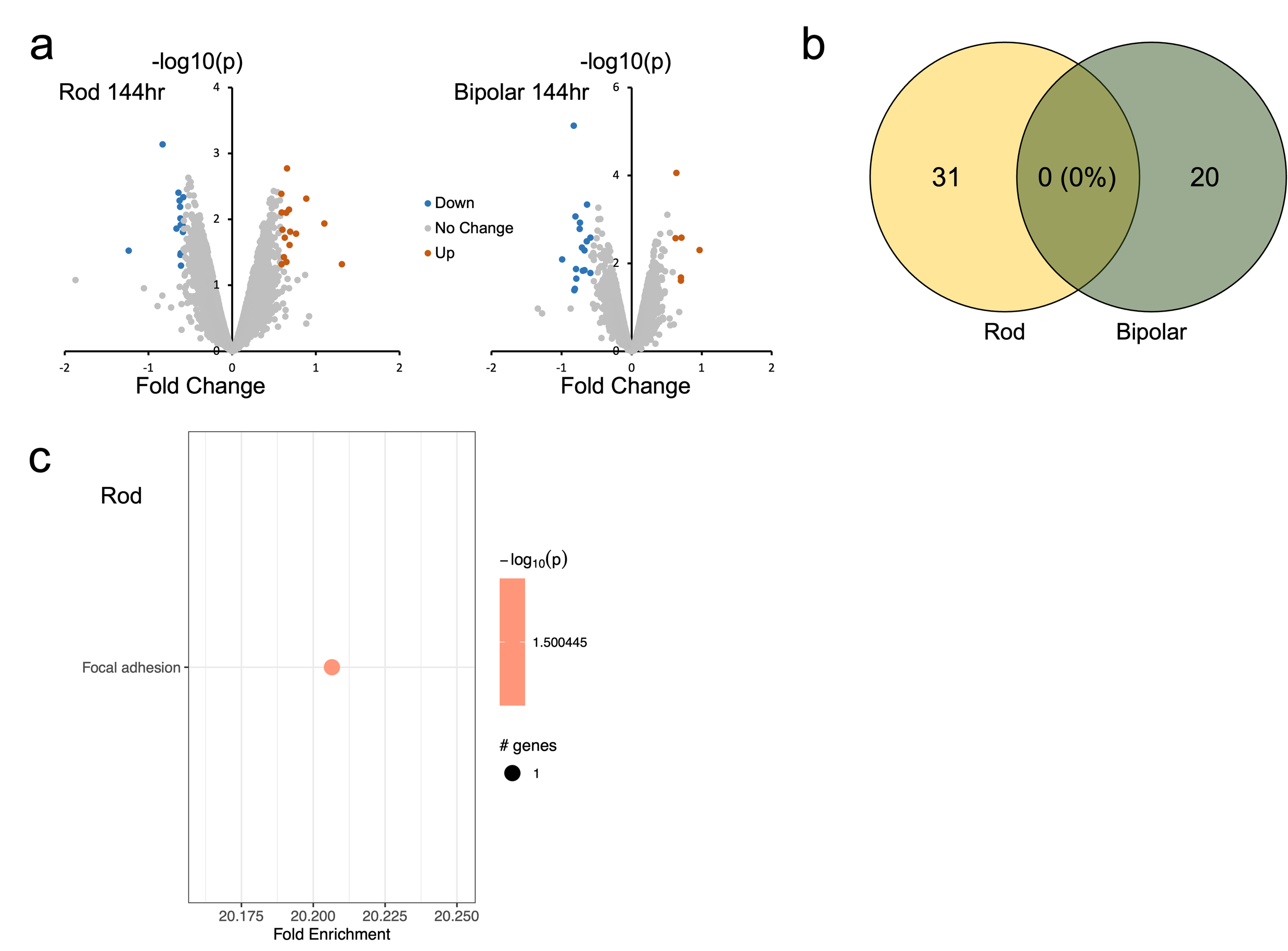

### Supp14.tif

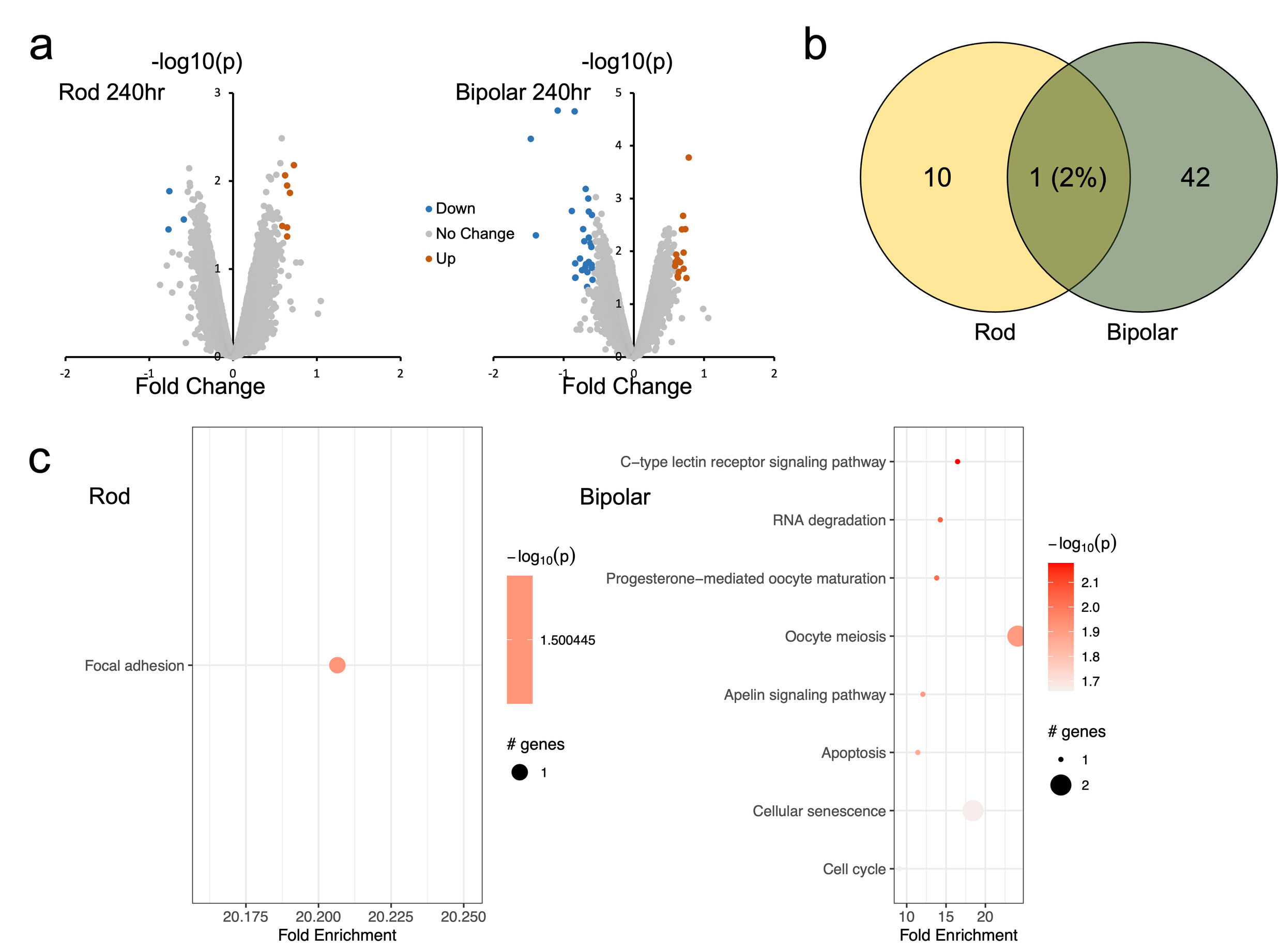

### Supp15new.tif

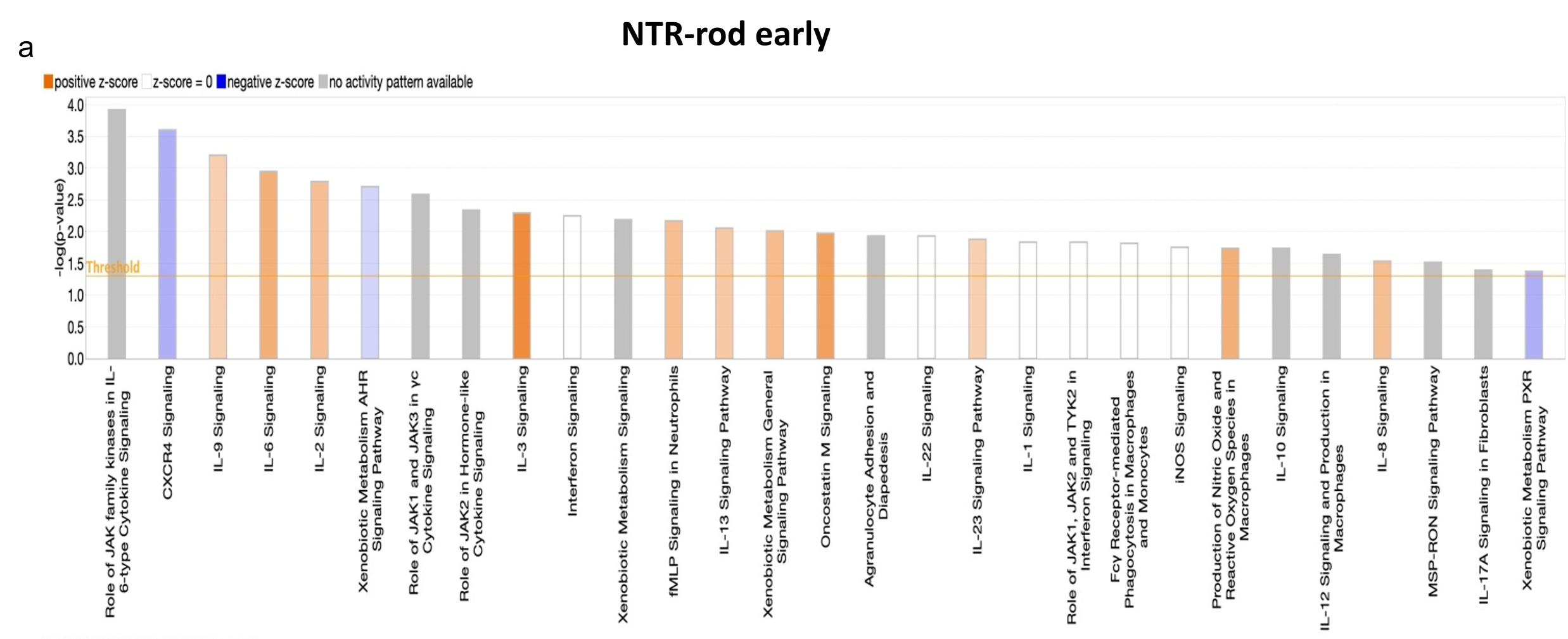

### Supp16new.tif

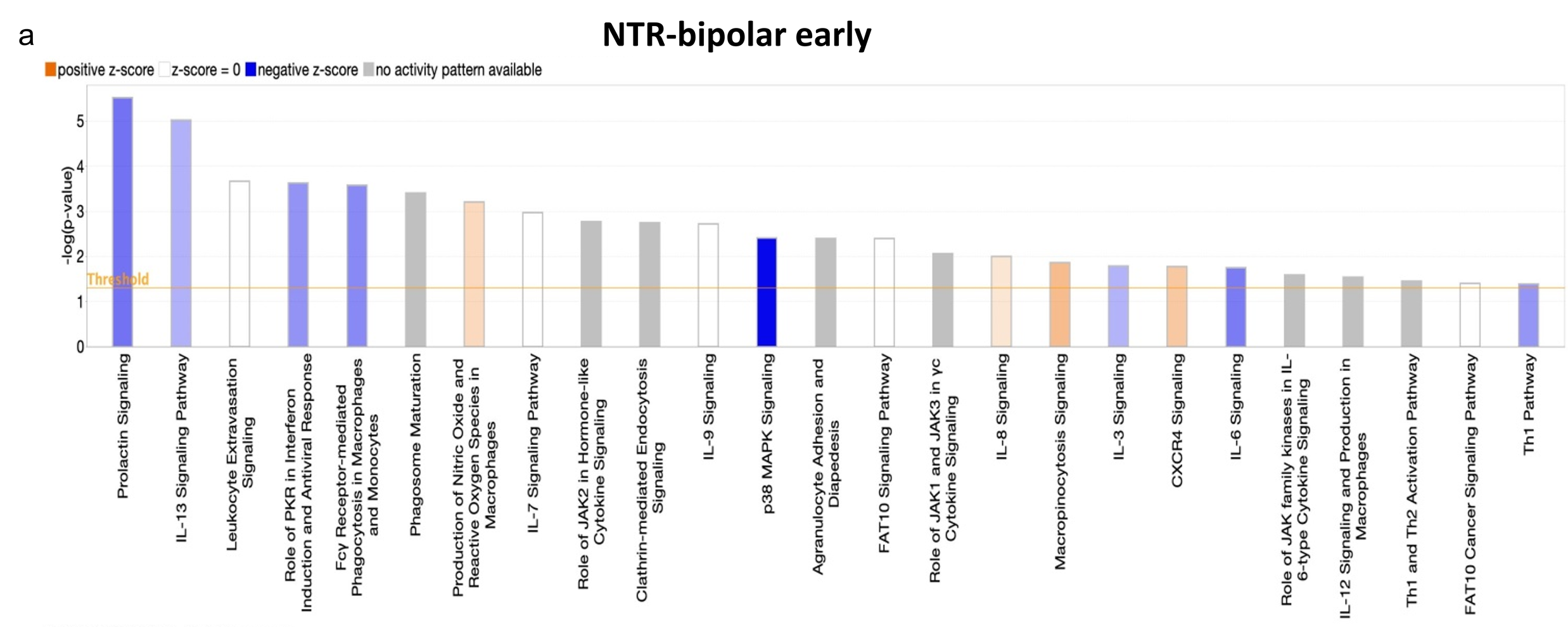

### Supp17new.tif

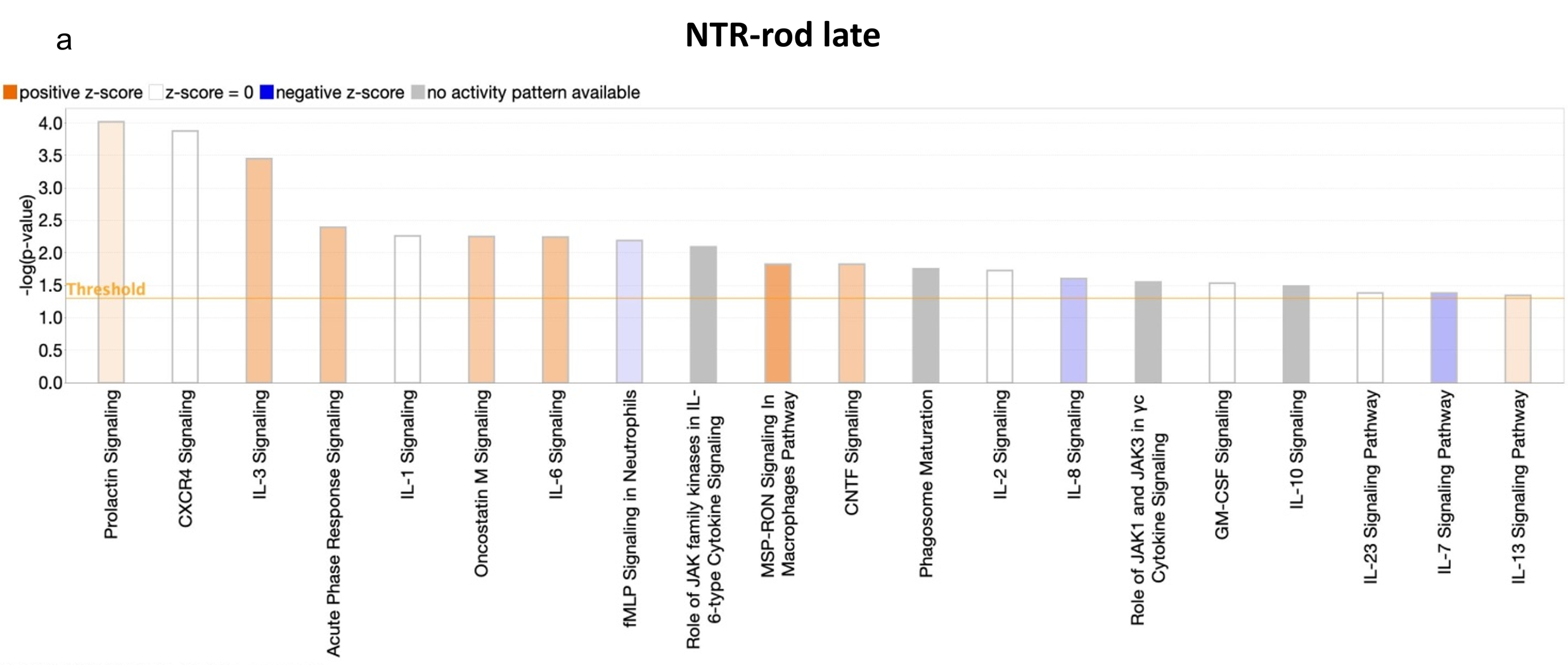

### Supp18new.tif

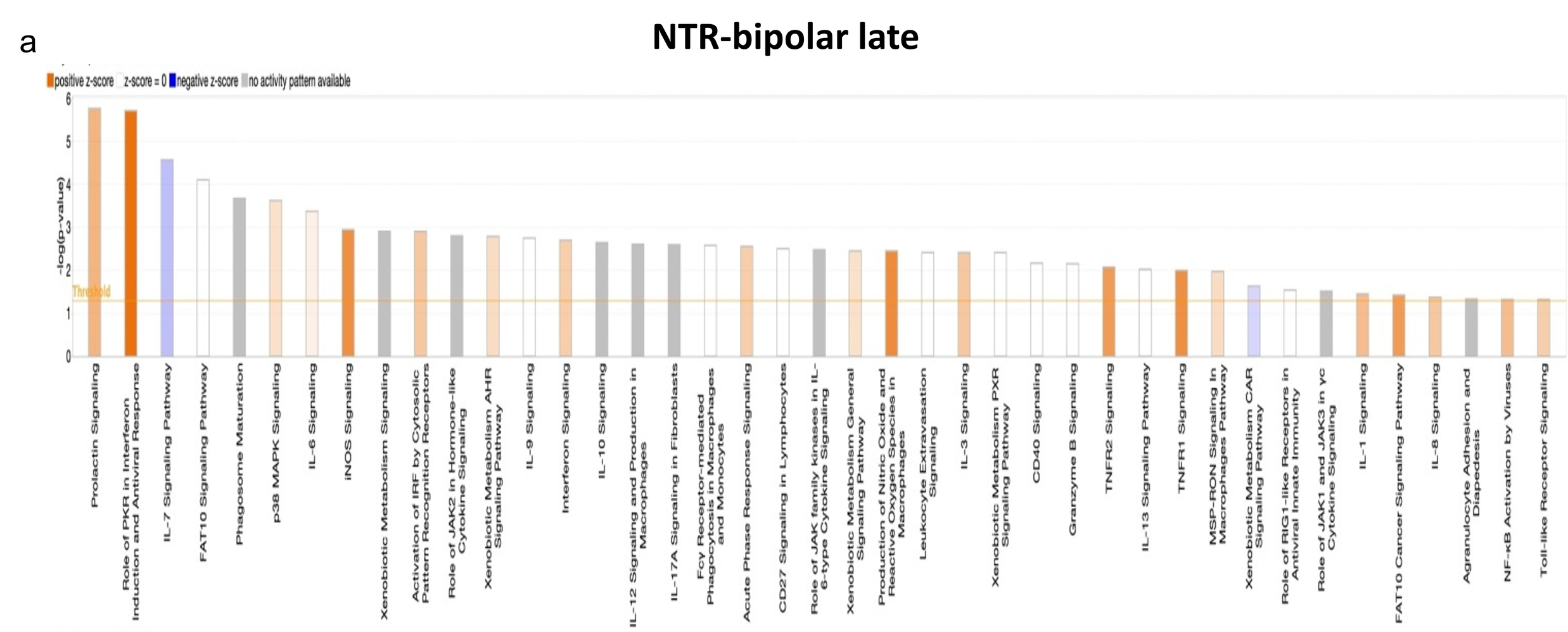

### Supp19.tif

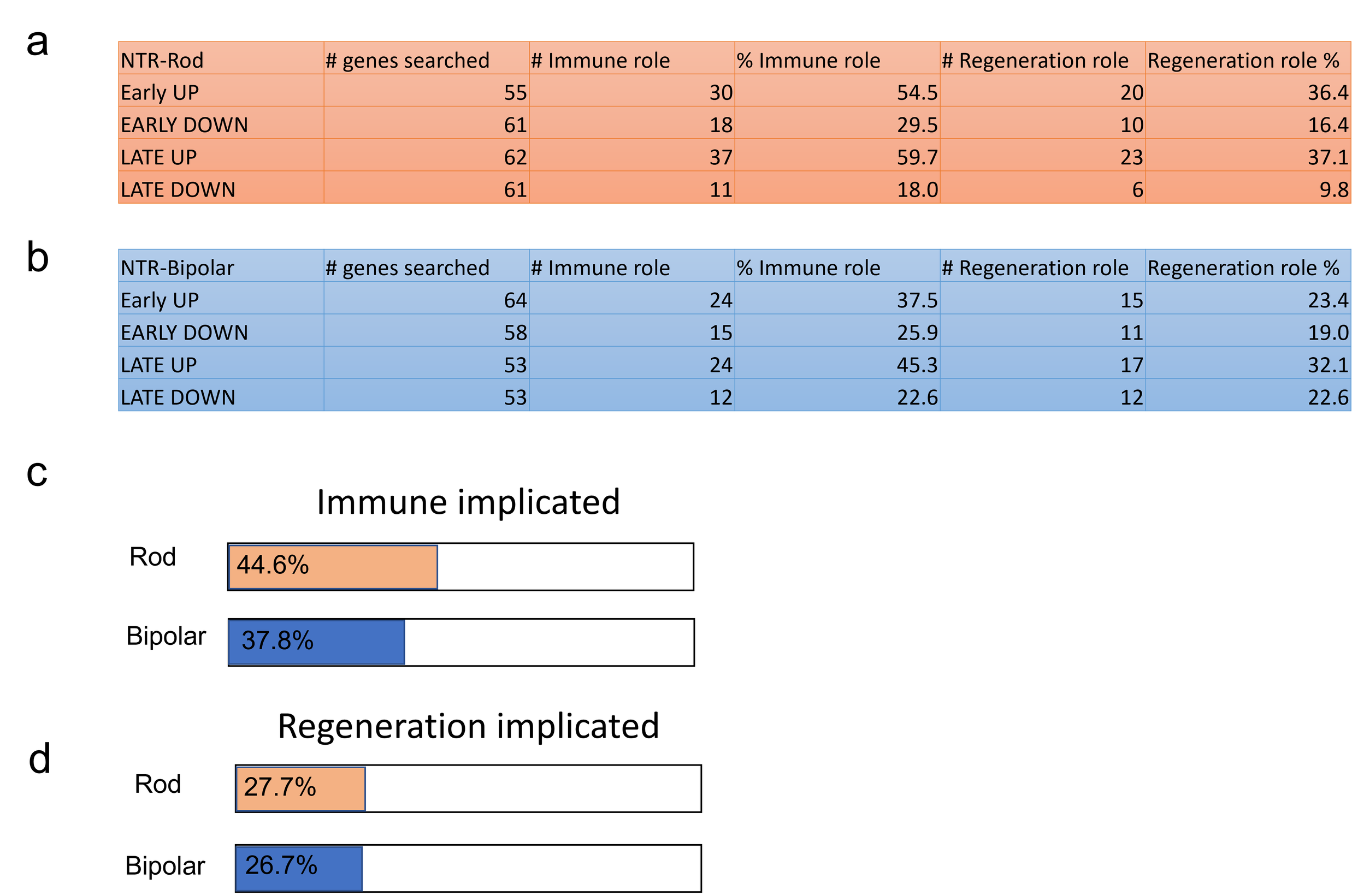

### SuppTable1.tif

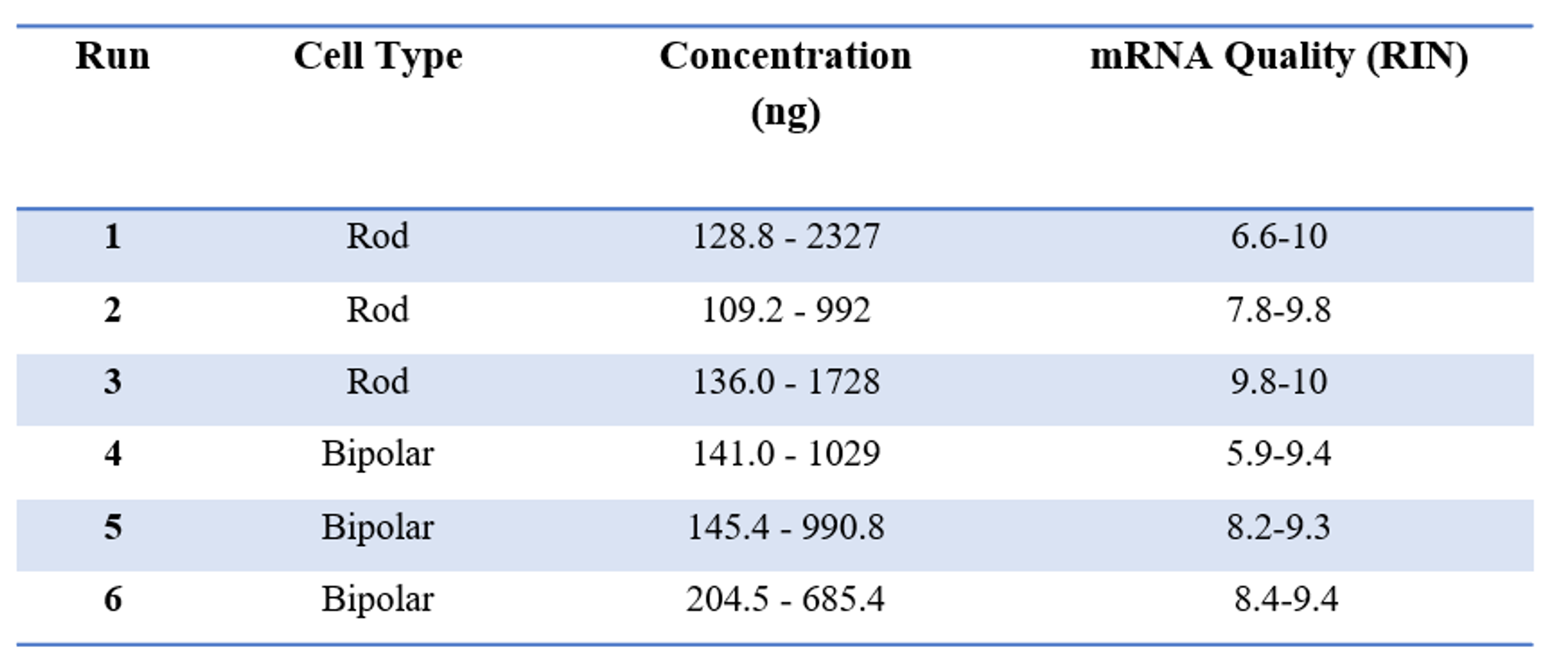

### SuppTable2.tif

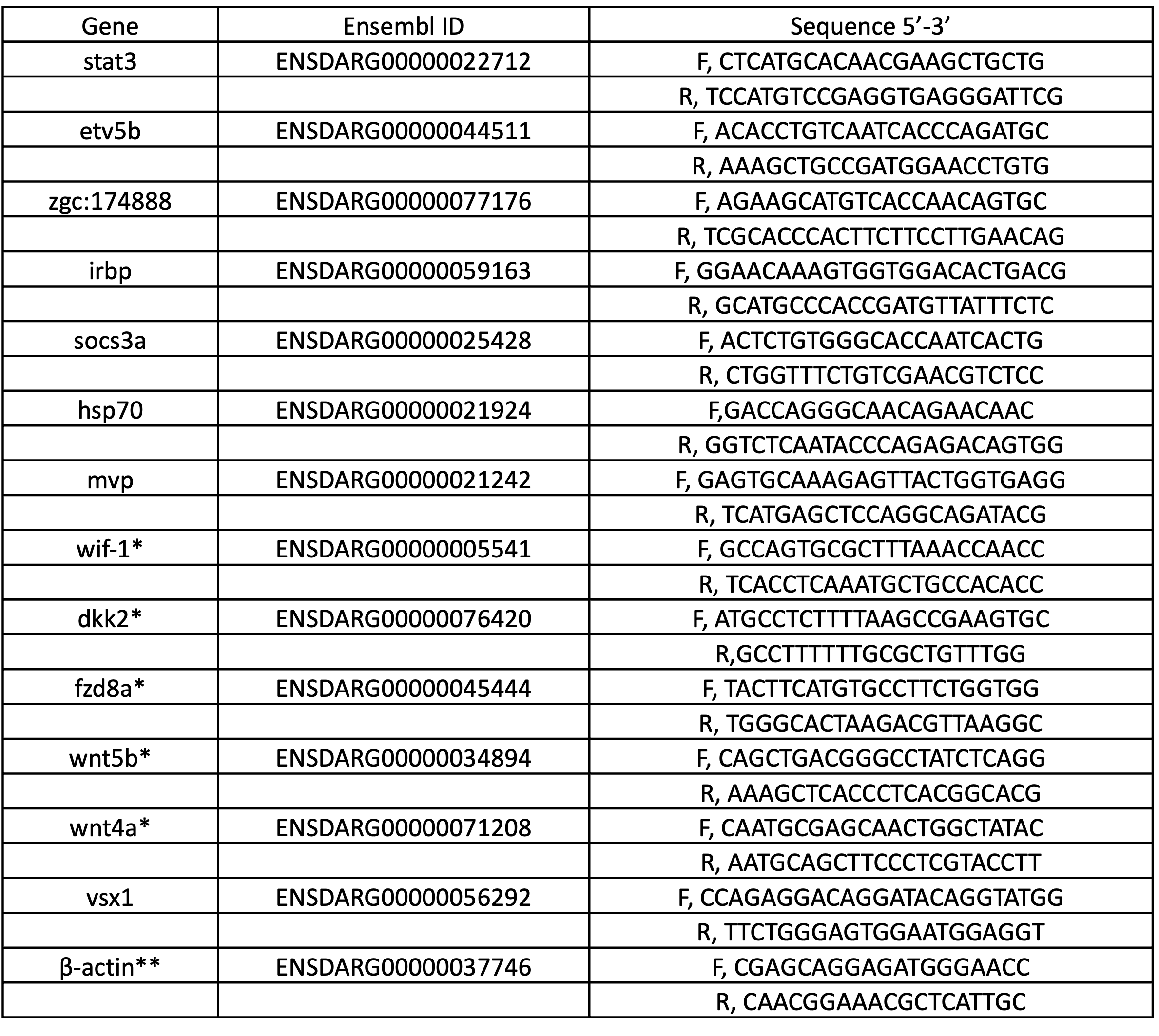
